## Supplementary Figure 1 for "HAYSTAC: A Bayesian framework for robust and rapid species identification in high-throughput sequencing data"

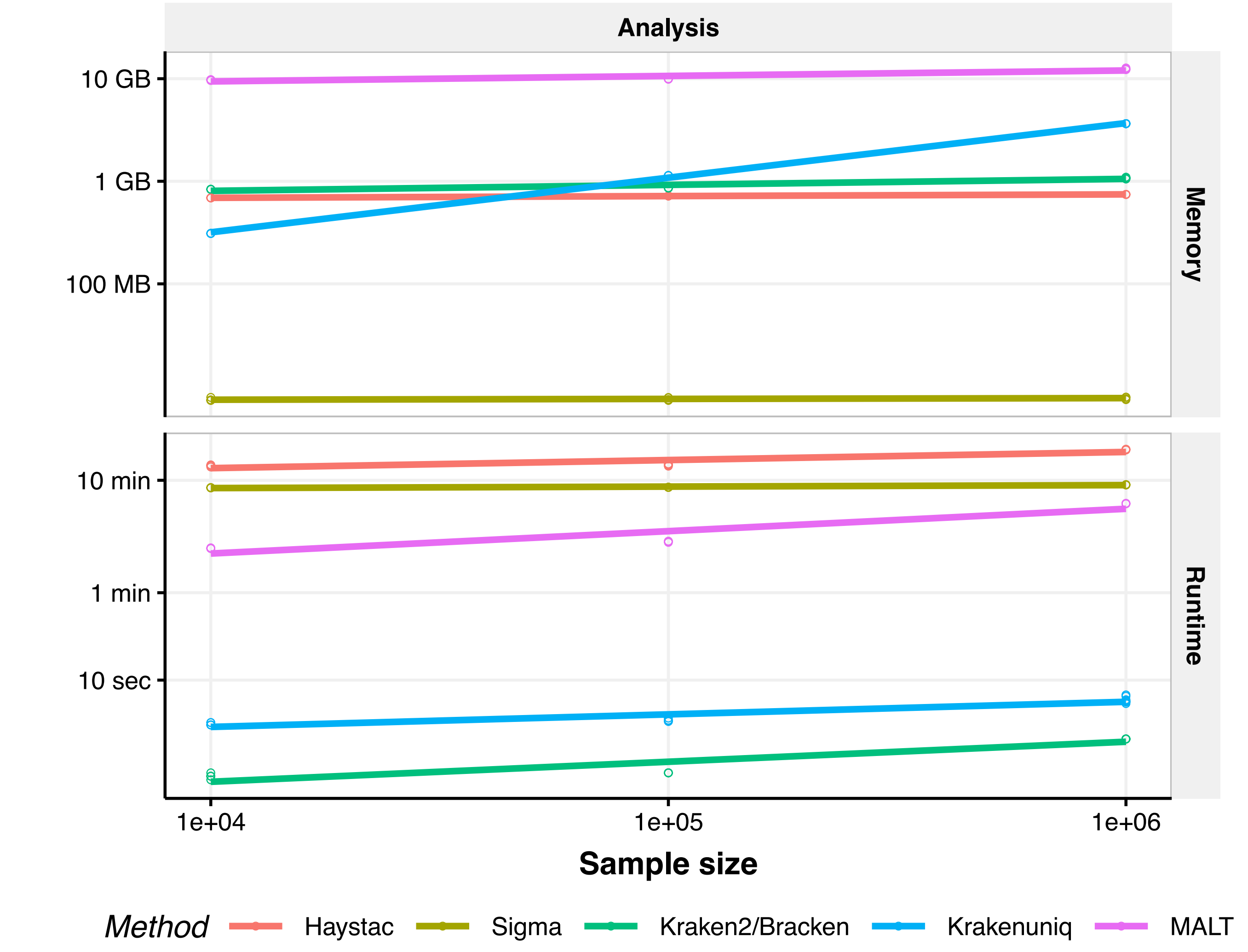


**Supplemental Figure 1.** Benchmarking for elapsed runtime and memory for HAYSTAC, Sigma, Kraken2/Bracken, Krakenuniq and MALT when analysing samples of $10$ K, $100$ K and $1$ M reads against a database of 500 species. Memory remains constant as sample size increases, and runtime in most methods (other than Kraken2/Bracken and Krakenuniq) scales with the database size rather than input sample size).
