## Supplementary Figure 2 for "HAYSTAC: A Bayesian framework for robust and rapid species identification in high-throughput sequencing data"

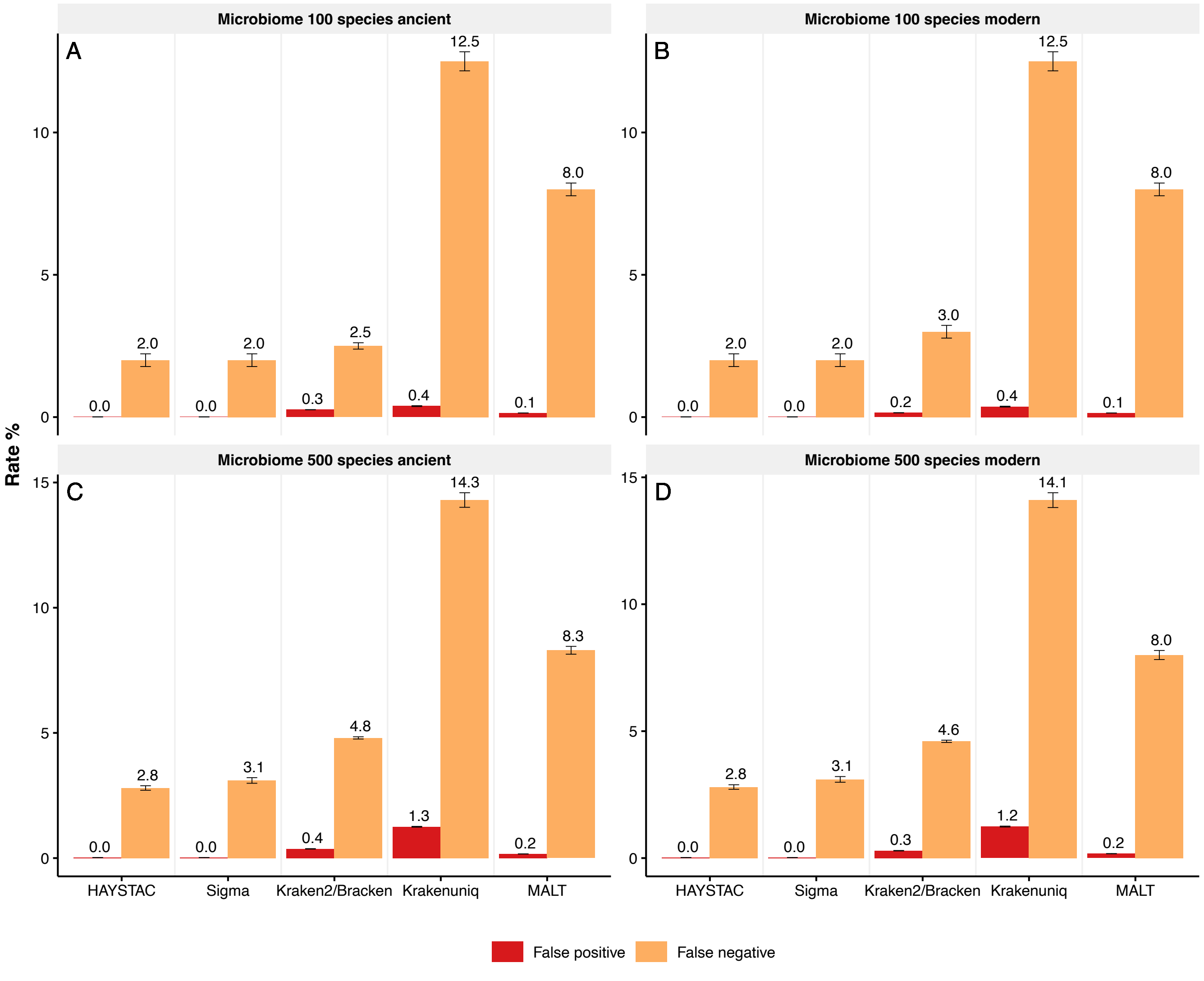


**Supplemental Figure 2.** False positive and negative rates per method for the simulated samples of the General Microbiome dataset.
