## Supplementary Figure 3 for "HAYSTAC: A Bayesian framework for robust and rapid species identification in high-throughput sequencing data"

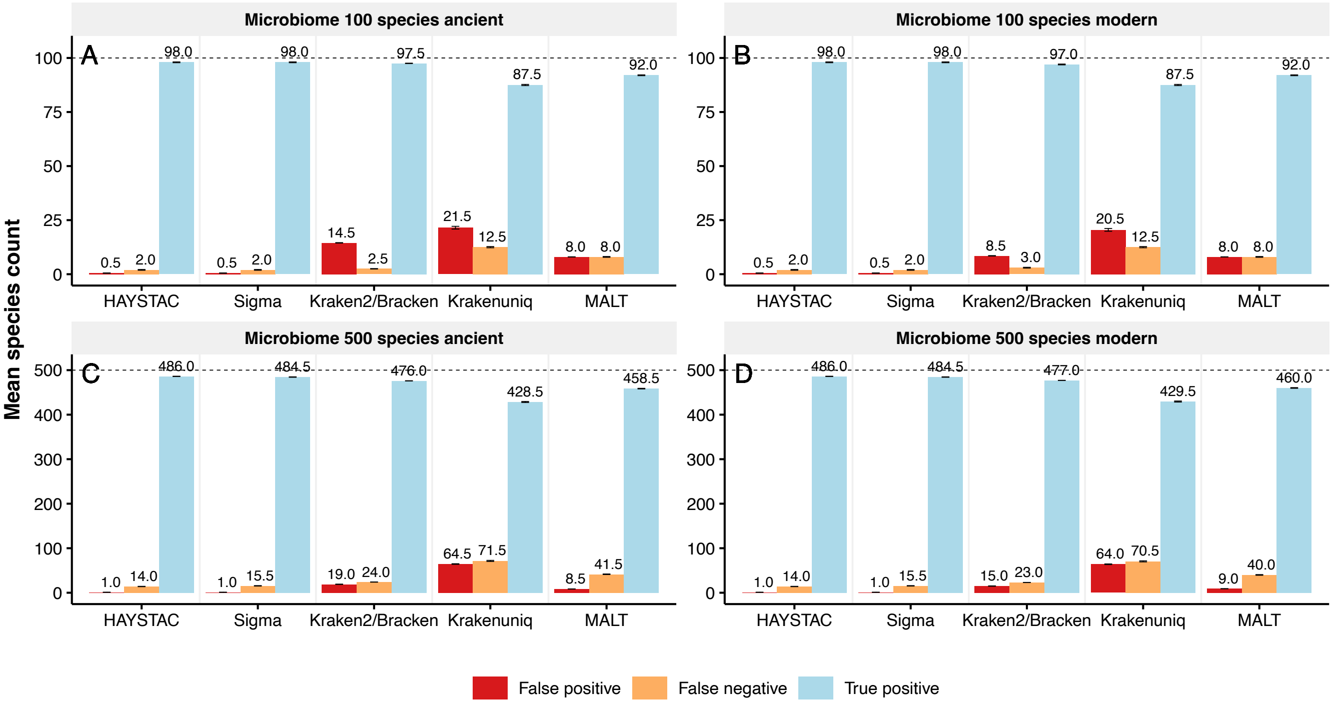


**Supplemental Figure 3.** Mean count of false positive (red), false negative (orange), and true detected species (blue) in the simulated General Microbiome dataset of 100 species ancient (n=2) (A), 100 species modern (n=2) (B), and 500 species ancient (n=2) (C) and 500 species modern (n=2) (D). The dotted line represents the average number of simulated species in each set of samples, and the numbers above the error bars represent the mean species count.
