## Supplementary Figure 4 for "HAYSTAC: A Bayesian framework for robust and rapid species identification in high-throughput sequencing data"

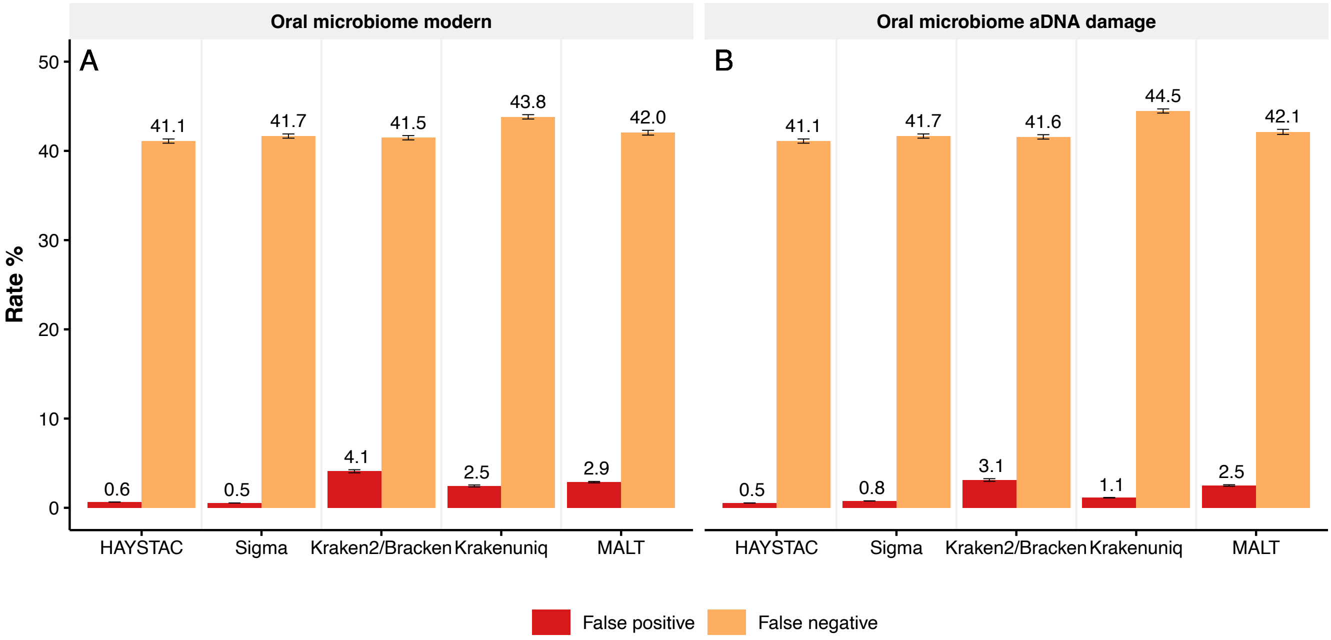


**Supplemental Figure 4.** False positive and negative rates per method for the simulated samples of the Oral Microbiome dataset.
