## Supplementary Figure 5 for "HAYSTAC: A Bayesian framework for robust and rapid species identification in high-throughput sequencing data"

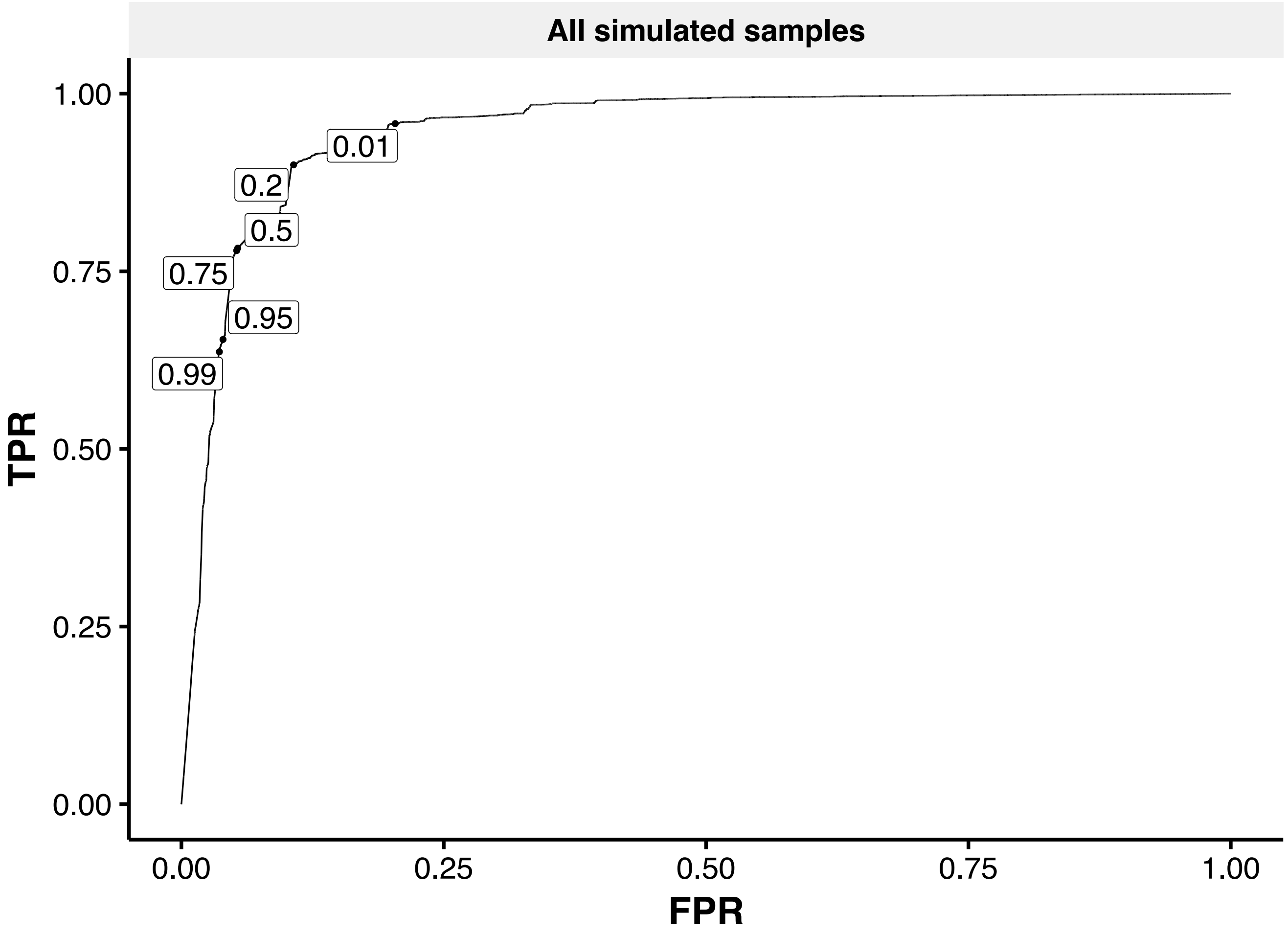


**Supplemental Figure 5.** Receiver operator curve analysis, showing the relationship between the true and false positive ratios (TPR and FPR respectively) to determine the default read posterior probability threshold for the Dirichlet assignment across all simulated samples. A rather conservative threshold of 0.75 is appropriate for most types of analysis. If the user requires a more relaxed threshold for discovery purposes the threshold can be lowered to 0.5. Respectively if a user is analysing deep sequencing data or wants to perform a more stringent identification the threshold can also be increased.
