## Supplementary Figure 6 for "HAYSTAC: A Bayesian framework for robust and rapid species identification in high-throughput sequencing data"

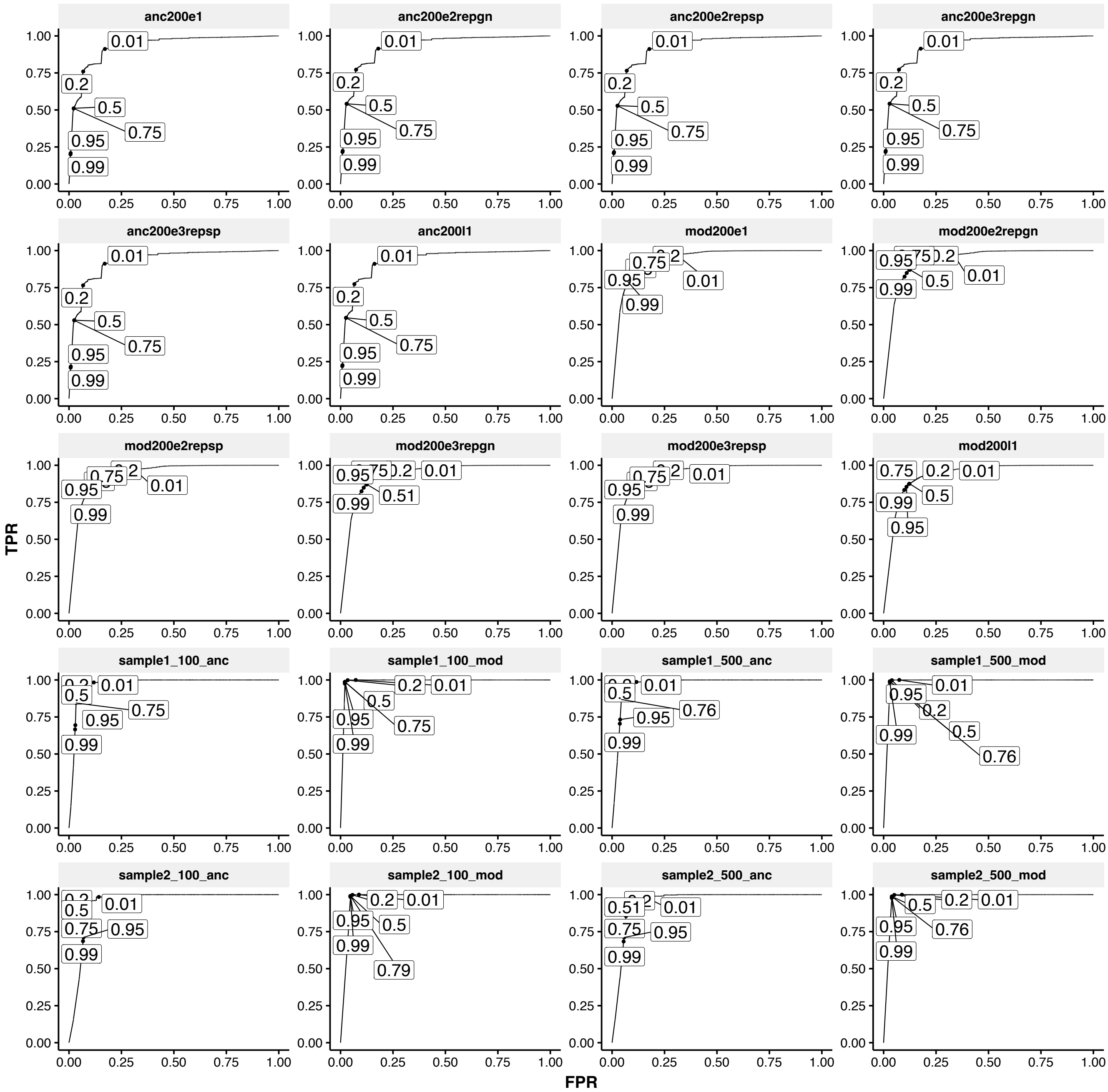


**Supplemental Figure 6.** Receiver operator curve analysis, showing the relationship between the true and false positive ratios (TPR and FPR respectively) to determine the default read posterior probability threshold for the Dirichlet assignment per sample. Here we can see how the read length and deamination levels affect the true positive rate. The user might want to consider these additional factors if they wish to change the default value of 0.75.
