## Supplementary Information for "HAYSTAC: A Bayesian framework for robust and rapid species identification in high-throughput sequencing data"

### HAYSTAC: Supplementary Information

#### Supplemental Appendix. Calculation of false positive, false negative and true positive rates

For the calculation of the false positive, false negative and true positive rates we used the following formulas:

$$FP_r = \frac{FP}{FP+TN} \tag{1}$$

$$FN_r = \frac{FN}{FN+TP} \tag{2}$$

$$TP_r = \frac{TP}{TP+FP} \tag{3}$$

where: $TP$ = number of true positives, $FP$ = number of false positives, $TN$ = number of true negatives, and $FN$ = number of false negatives. For a true positive identification we require an abundance equal or higher than $0.01\%$.
